# A steady state pool of calcium-dependent actin is maintained by Homer and controls epithelial mechanosensation

**DOI:** 10.1101/2025.09.05.674388

**Authors:** Kenji Matsuzawa, Makoto Suzuki, Yuma Cho, Ryoya Fujinaga, Junichi Ikenouchi

## Abstract

Epithelial cells are inherently contractile and in homeostasis, tissue integrity is maintained by balancing the uneven contractile forces in neighboring cells at the cell-cell interface. By contrast, epithelial cells can utilize an imbalance in contractile force to communicate various information to induce tissue-wide response as in wound healing. Contractility is generated and processed at the apical junctional complex (AJC) by the dynamic behavior of the actin cytoskeleton. Calcium signaling can pattern cellular responses based on its reach and amplitude and the actin cytoskeleton is supported by its wide ranging effects on actin regulators. Calcium transients regulate various cell behaviors associated with actin remodeling, such as in damage response and developmental morphogenesis. Here we report that calcium maintains an adaptive pool of AJC-associated actin that is sensitive to tension and encoded by calcium dynamics. For this, the recently identified epithelial polarity module Homer-MUPP1/PatJ is required. Homer regulates calcium signaling in various tissue contexts through interaction with numerous components of the endoplasmic reticulum (ER) and plasma membrane (PM) calcium signal toolkit. Knockout of either Homer or MUPP1/PatJ attenuated tension-induced calcium response and severely disrupted wound healing migration, which is dependent on guidance input through AJC tension. We also show that Homer is integral to early embryonic neurodevelopment as its suppression causes failure of neural tube closure. Our findings highlight the critical role of localized calcium dynamics on AJC actin remodeling and cellular behavior, elucidating the means of tissue coordination through intercellular tension.

**Significance statement:** This study uncovers a novel mechanism by which localized calcium dynamics at the apical junctional complex maintain an adaptive, tension sensitive pool of actin, regulated by the epithelial polarity scaffolds Homer and MUPP1/PatJ. Importantly, this mechanism operates without perturbing epithelial polarity, indicating a specific means to modulate tissue mechanics. By linking mechanical forces to localized calcium amplification, this module enables precise mechanosensation, coordinating collective behaviors such as epithelial wound healing and neural tube closure in *Xenopus*. These findings redefine our understanding of intercellular tension sensing in epithelial tissues and highlight the Homer–calcium signaling axis as a key driver of tissue morphogenesis and homeostasis, with far reaching implications for developmental biology, regenerative medicine, and neural tube defect pathogenesis.

## Main text

### Calcium maintains a steady state pool of AJC-associated actin and its dynamics underlies AJC deformation

Contractility at epithelial adhesions is balanced in homeostasis^1,2^ and conveys mechanical information during coordinated tissue response^3,4^. Calcium signaling also regulates a wide range of collective behavior^5–11^ through its effect on the actin cytoskeleton. We wondered what role, if any, calcium might play in regulating steady state epithelial cell dynamics. We first examined calcium dynamics at the apical junctional complex (AJC) by expressing the membrane-tethered calcium indicator GCaMP-CAAX in the mouse mammary epithelial cell EpH4 (Figure 1A). While there were occasional bursts, the signal by and large held steady. Kymographs drawn perpendicular to the AJC plane to visualize junction movement, i.e. the tug of war between cells, a proxy for intercellular tension, with the GCaMP signal appeared to show a slight tendency towards higher signal intensity during more labile periods (Figure 1B). We therefore analyzed the hierarchical relationship between junction movement and calcium dynamics by cross-correlation analysis, comparing the lags obtained from the experimental dataset to a control dataset consisting of randomized pairs of the experimental junction movements and calcium dynamics. While we found no significant difference in the means (control = –63 sec and experimental = –38 sec with the negative lag indicating that changes in calcium dynamics follows junction deformation), the experimental lags were significantly less varied and tightly clustered, suggesting that there is a meaningful relationship between junction deformation and calcium dynamics (Figure S1A).

**Figure 1.**
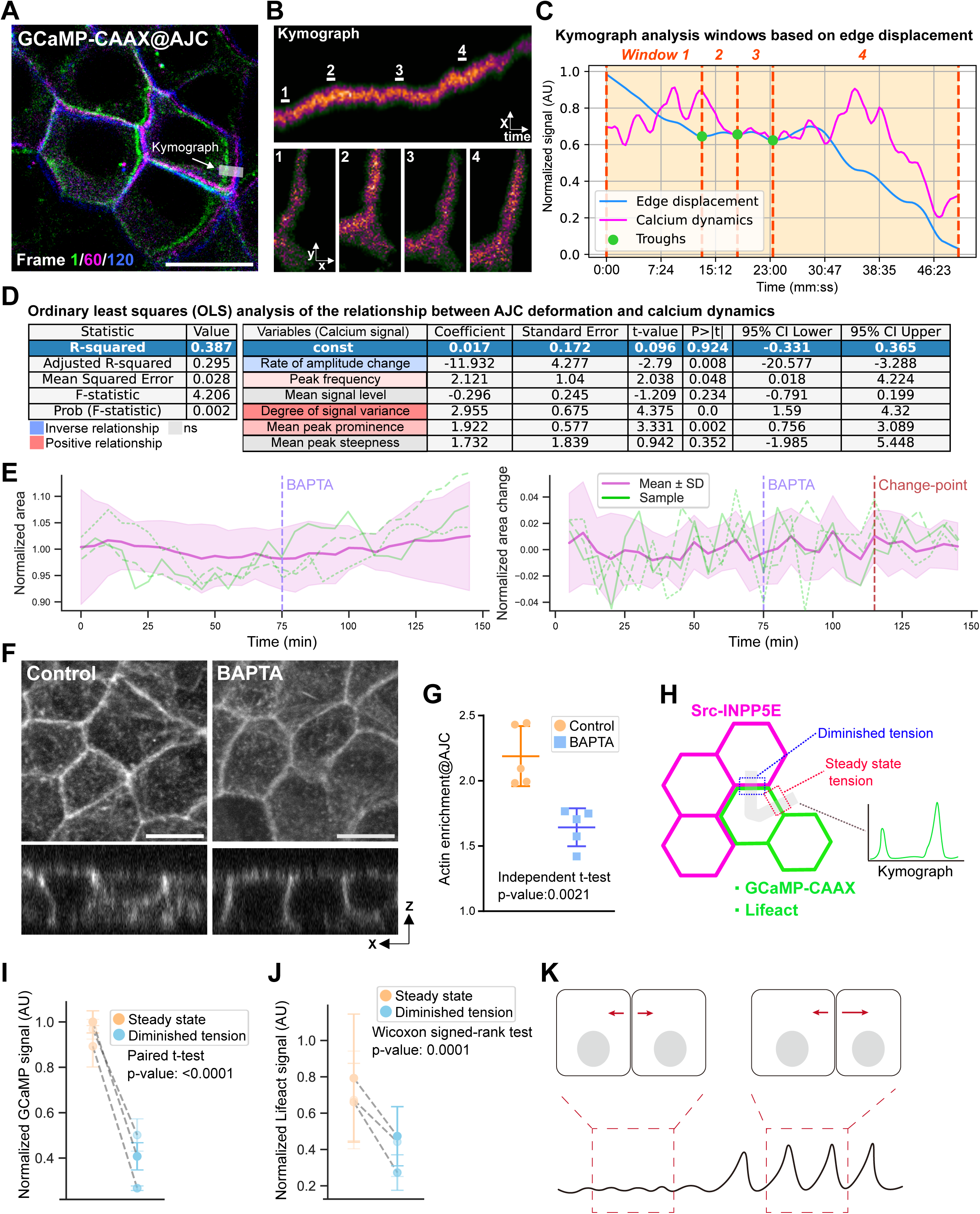
Calcium maintains a steady state pool of AJC-associated actin and its dynamics underlies AJC deformation. (A) Time overlay of the GCaMP signals at AJC from a representative time lapse image of EpH4 cells expressing GCaMP-CAAX. The shaded area shows the kymograph region of interest analyzed in (B) and (C). Scale bar = 20 µm. (B) Kymograph and time projection images (average intensity) of four frames at the indicated time points. (C) Edge movement and corresponding changes in GCaMP signal were derived from processed kymograph, expressed as absolute values of displacements. The interval between troughs of the movement data was used to set up analysis windows that roughly reflect the tendency of edge movement. Cumulative GCaMP signal change and the number of GCaMP peaks were determined for each analysis window. All data were normalized for the duration of the analysis window and any window containing a zero value for any variable were thrown out. A total of 94 analysis windows were obtained from 12 cell junctions obtained over three independent experiments and 6 cells. (D) Ordinary least squares analysis of the kymograph data. The analyzed variables were as follows: ‘Rate of amplitude change’, which describes the slope of the signal; ‘Peak frequency’, which is the number of peaks per window, ‘Mean signal level’; ‘Degree of signal variance’, which describes how much the signal varies within the window; ‘Mean peak prominence’, which is how high the peak is over baseline; and ‘Mean peak steepness’, which is how sharply the peaks rise. (E) Cell areas were quantified from EpH4 cells expressing the TJ marker GFP-Claudin-3 to visualize the AJC outline. The timings of BAPTA treatment and change point appearance are indicated. Means and standard deviation of 12 analyzed junctions are shown with three representative individual cell traces overlaid. (F) Phalloidin staining of EpH4 cells treated with BAPTA-AM for 30 min. Scale bar = 20 µm. (G) Quantification of (F). ZO-1 and E-cadherin staining were used to define AJC and lateral regions to quantify actin enrichment at AJC. Mean and standard deviation are indicated. Individual dots are means of one field of view and five independent images obtained over three independent experiments were analyzed. Also see Figure S2B. (H) Schematic of the experiments artificial manipulating intercellular tension. GCaMP-CAAX– or EGFP-Lifeact-expressing cells were co-cultured with cells stably expressing the lipid phosphatase INPP5E, which reduces PM anchoring of F-actin via Ezrin by depleting PI(4,5)P2, thereby reliably creating cell junctions subject to uneven tension within the same cell. (I) and (J) Quantification of calcium dynamics (I) and actin enrichment (J). Signal data was normalized to the control, steady state cell junction not subject the INPP5E-expressing cells. Individual means and standard deviations of three independent experiments with 12 cell junctions tested per experiment are shown with the control and experimental pairs indicated by dashed lines. (I) Schematic of the relationship between intercellular tension and calcium dynamics.

In order to understand what aspects of the calcium dynamics is relevant to AJC deformation, each kymograph time series was split into analysis windows based on magnitude of junctional displacement and corresponding variables reflecting signal shape and intensities were derived (Figure 1C). The GCaMP signal was pulsatile, reflecting the influx-uptake cycle of calcium dynamics. Plotting the signal intensity and shape variables against edge displacement revealed only a weak linear relationship for the variance in signal intensity (Figure S1B). We then performed ordinary least squares (OLS) analysis, which allows examination of the relationship between multiple explanatory variables and a single outcome variable. We found that calcium dynamics as a whole significantly accounts for AJC deformation, capable of explaining nearly 40% of the variability in junction movement (Figure 1D). Of the calcium signal aspects, the most significant predictor was the degree of variance in signal intensity, showing a positive relationship with edge displacement. Peak frequency and prominence were also positively associated; meanwhile, the rate of change in signal intensity showed an inverse relationship. The positively associated variables all describe variance in calcium levels, which we identify here as a strong correlative factor of sustained AJC deformation, with greater fluctuation being predictive of greater edge displacement. Appropriately, suppression of cytoplasmic calcium by the cell permeable calcium chelator BAPTA-AM diminished AJC deformation, as indicated by analysis of cell area fluctuation (Figure 1E). Change point detection is a method to identify when statistically significant changes occur in a time series. Applying this to the rate of cell area change time series, we identified a unique change point following BAPTA treatment (Figure 1E). Re-analyzing the dataset pre– and post-change point revealed that cell area increased and the variability in cell malleability trended downward post-change point (Figure S1C). Since actin contraction underpins AJC deformation, we expected that calcium suppression would affect AJC-associated actin architecture. BAPTA-treated cell lacked the taut appearance of control cells, suggesting that intercellular tension is diminished. There was a marked decrease in F-actin enrichment at AJC under calcium chelation indicating that the AJC F-actin population is susceptible to decreased cytoplasmic calcium levels (Figures 1F and 1G). Taken together, these observations suggest that calcium maintains a pool of AJC-associated F-actin that is sensitive to—and feeds back to—intercellular tension.

To further test the idea that tension-sensitive calcium regulates AJC actin level, we subjected GCaMP-CAAX– and EGFP-Lifeact-expressing cells to differential mechanical input at cell junctions of single cells. To do so, we established cells stably expressing a membrane-targeted lipid phosphatase INPP5E to downregulate PI(4,5)P2, which downregulates plasma membrane (PM)-anchored F-actin and consequently decreases junctional tension^24^. When these cells are co-cultured with wild type (WT) cells expressing the biosensors, the junction touching the INPP5E-expressing cell should experience diminished tension compared to non-contacting junctions; the sensors were expressed mosaically to ensure there was no signal overlap from an adjacent sensor-expressing cell (Figure 1H). We then analyzed signals at both junction types simultaneously by a kymograph, which revealed that both baseline calcium level and F-actin enrichment were suppressed at junctions neighboring the INPP5E-expressing cell (Figures 1I and 1J).

Thus far, we have shown that calcium dynamics at AJC is enhanced during periods of greater junction deformation relative to quiescent periods, leading to greater variance in calcium dynamics, and that suppression of cytoplasmic calcium diminishes AJC deformation and F-actin enrichment. As a visual output of intercellular tension, greater AJC deformation is suggestive of imbalance in intercellular tension and lack of movement of an equilibrium state in the contractility between cells. The magnitude of variance in calcium dynamics then is potentially a readout of the difference in contractility at AJC. While it is yet unclear as to what element of AJC deformation calcium is actually sensitive to—fluctuations in membrane tension or actomyosin contraction for example—these results suggest that calcium dynamics could be a way for cells to sense and transduce the disparity in mechanical input by decoding it into the fluid pool of calcium-regulated AJC actin to increase the opposing contractility in response (Figure 1K).

### Calcium-dependent AJC actin is regulated by Homer-MUPP1/PatJ

Recently, the Homer family of neuronal scaffolding proteins were identified as accessory components of the epithelial polarity machinery^4^. Homer localizes to the dendrite^25^, the postsynaptic density^5,13^ and the growth cone^12^ in neurons where they interact with, and often couple, resident calcium channels of the PM and ER to control actin-dependent processes such as synaptic maturation and growth cone turning; Homer itself was also reported to possess actin bundling activity^26^. Additionally, Homer links the IP3 receptor to glutamate receptors in the postsynapse^7^ and suppressing Homer severely attenuated Store-operated calcium entry (SOCE) in platelets^14^ and vascular smooth muscle cells^6^, making it an attractive candidate to regulate calcium dynamics at AJC (Figure 2A). We confirmed the localization of the two expressed Homer isoforms (1 and 3) at the AJC of EpH4 cells and established knockout cells. (Homer dKO, Figures 2A and 2B). Using the localization of E-cadherin (adherens junction, AJ) and claudin-4 (tight junction, TJ) as markers, we found that loss of Homer did not visibly perturb epithelial polarity (Figure 2C). However, we noted that Homer dKO cells were notably diminished in height compared to either wild type (WT) or Homer3-rescued cells (Homer3res) in co-culture. Examination of cortical actin distribution revealed that AJC enrichment in Homer dKO cells was compromised as in BAPTA-treated cells, suggesting that loss of AJC actin causes diminished cell height in Homer deficient cells due to a reduction in apical tension (Figure 2D). Likewise, AJC deformation dynamics was significantly diminished, similar to WT cells post-calcium chelation, as indicated by the decrease in the rate of cell area change (Figure 2E).

**Figure 2.**
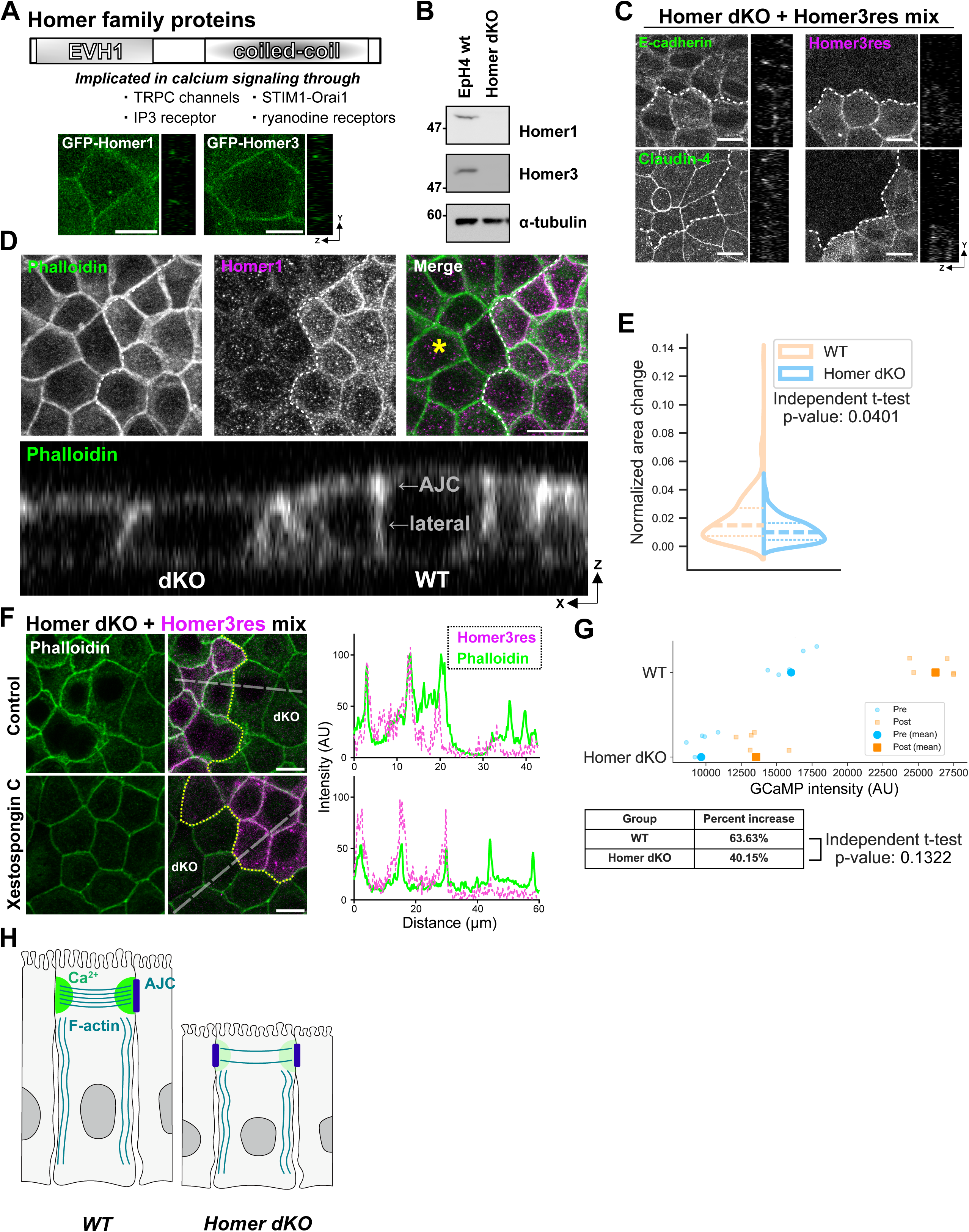
Calcium-dependent AJC actin is regulated by Homer. (A) Domain structure of Homer family proteins. Known interactors in calcium signaling are annotated. Images of GFP-Homers expressed in EpH4 cells are shown below. Scale bar = 10 μm. (B) Whole-cell lysates of WT and Homer dKO cells were immunoblotted with the indicated antibodies. (C) Immunofluorescence images of Homer dKO cells and Homer dKO cells expressing CRISPR resistant mScarlet-Homer3 stained for the TJ marker Claudin-4 (top panels) and the AJ marker E-cadherin (bottom panels). Scale bar = 10 µm. (D) Phalloidin staining of a co-culture of WT and Homer dKO cells. Scale bar = 20 µm. (E) The rate of cell area change was quantified from WT and Homer dKO cells expressing GFP-Claudin-3. At least 40 cells were quantified over three independently acquired time lapse experiments. Violin plot shows the median with 25^th^ and 75^th^ percentiles. (F) Phalloidin staining of a co-culture of Homer dKO and Homer dKO cells expressing CRISPR resistant mScarlet-Homer3. Cells were treated with Xestospongin C for 30 min. Line scans are shown at right. Scale bar = 10 µm. (G) Response of WT and Homer dKO cells exposed to the calcium ionophore A23187. Discrete points indicating mean fluorescence of 5 time points before and after ionophore treatment from a single time series are shown with the means. (H) Schematic showing the effect of depressed calcium on AJC actin in Homer dKO cells.

The endoplasmic reticulum (ER) is a major calcium store that releases calcium via IP3 and ryanodine receptors (IP3R and RyR). Given previous reports, we asked whether Homer regulates AJC actin through ER calcium release. We found that the difference in baseline AJC actin level between WT/Homer3res and Homer dKO cells could be erased by inhibiting calcium release through IP3 receptor (Figure 2F). We sought to examine calcium dynamics in Homer dKO cells but GCaMP-CAAX signal was barely perceptible in a large proportion of cells under normal culture conditions over repeated attempts to establish stable cell lines. The established Homer dKO cells, however, were responsive to stimulation with the calcium ionophore, which suggests that baseline calcium level is severely attenuated in Homer dKO cells (Figure 2G). Thus, calcium and Homer controls the labile, tension-sensitive pool of AJC actin, which is dispensable for the formation epithelial adhesions (Figure 2H).

Homer was recently shown to biochemically interact with the Crumbs complex component PatJ (Pals1-associated tight junction protein)^4^. We thus asked whether PatJ is involved in Homer localization at AJC. We confirmed that both Homers1 and 3 bind PatJ and its homolog MUPP1 (Multi-PDZ domain protein 1) by immunoprecipitation (Figure S3A). Homer is composed of the N-terminal EVH1 domain that mediates protein-protein interactions and the C-terminal coiled coil region, through which it oligomerizes^27,28^. Examining the Homer-MUPP1/PatJ interaction further, we found that the EVH1 domain is sufficient to bind PatJ (Figures S3B and S3C). Establishing MUPP1/PatJ knockout (M/P dKO) cells revealed that the correct localization of both Homer isoforms is dependent on MUPP1/PatJ (Figures S3D and S3E). Surprisingly, although PATJ and MUPP1 are considered essential for TJ formation through their interactions with various TJ components, including ZO proteins, cells lacking both PATJ and MUPP1 showed no significant alterations in the localization of key TJ and AJ components, aside from the loss of Homer localization (Figure S3F). We conclude that Homer-MUPP1/PatJ regulates AJC actin through a calcium-dependent mechanism but is dispensable for epithelial junction formation.

### Signaling through Homer-MUPP1/PatJ controls mechanosensation in cultured epithelial cells

We have shown so far that localized intracellular calcium maintains a pool of actin at AJC that is sensitive to steady state levels of intercellular tension, which requires Homer-MUPP1/PatJ. Variously described as transients, sparks and flashes, elevation of cytoplasmic calcium has been observed in epithelial tissue in association with acute mechanical stimuli during junction damage repair, tissue wound response and developmental morphogenesis^1–3,20,22,29–32^. Curious as to whether Homer-MUPP1/PatJ in fact mediates cellular response to steady state levels of mechanical perturbation, we stably expressed a nucleus-targeted GCaMP (NLS-GCaMP) in WT, Homer dKO and MUPP1/PatJ dKO cells in order to observe whole cell response. Since Homer and MUPP1/PatJ dKO cells cannot assemble the mechanosensitive calcium-dependent pool of AJC actin, we would expect them to show a diminished calcium response to tensile stress. Cells were co-cultured with WT cells that express a blue light-induced activator of RhoA GTPase (OptoRhoGEF) in order to apply tension at will^33^. In WT cells, irradiation with blue light evoked a sharp rise in the GCaMP signal that was sustained for minutes after stimulation. By contrast, the signal quickly dissipated to baseline levels after the initial elevation in both Homer and MUPP1/PatJ dKO cells (Figures 3A and S4A). Cluster analysis revealed three response classes: cluster 1 cells failed to sustain the response, cluster 2 cells showed an exaggerated response and cluster 3 cells maintained an elevated response. Responses of WT cells were predominantly grouped in clusters 2 and 3 while those of Homer and MUPP1/PatJ dKO cells were almost exclusively found in cluster 1, indicating that mechanosensation is deficient in these cells (Figure S4B).

**Figure 3.**
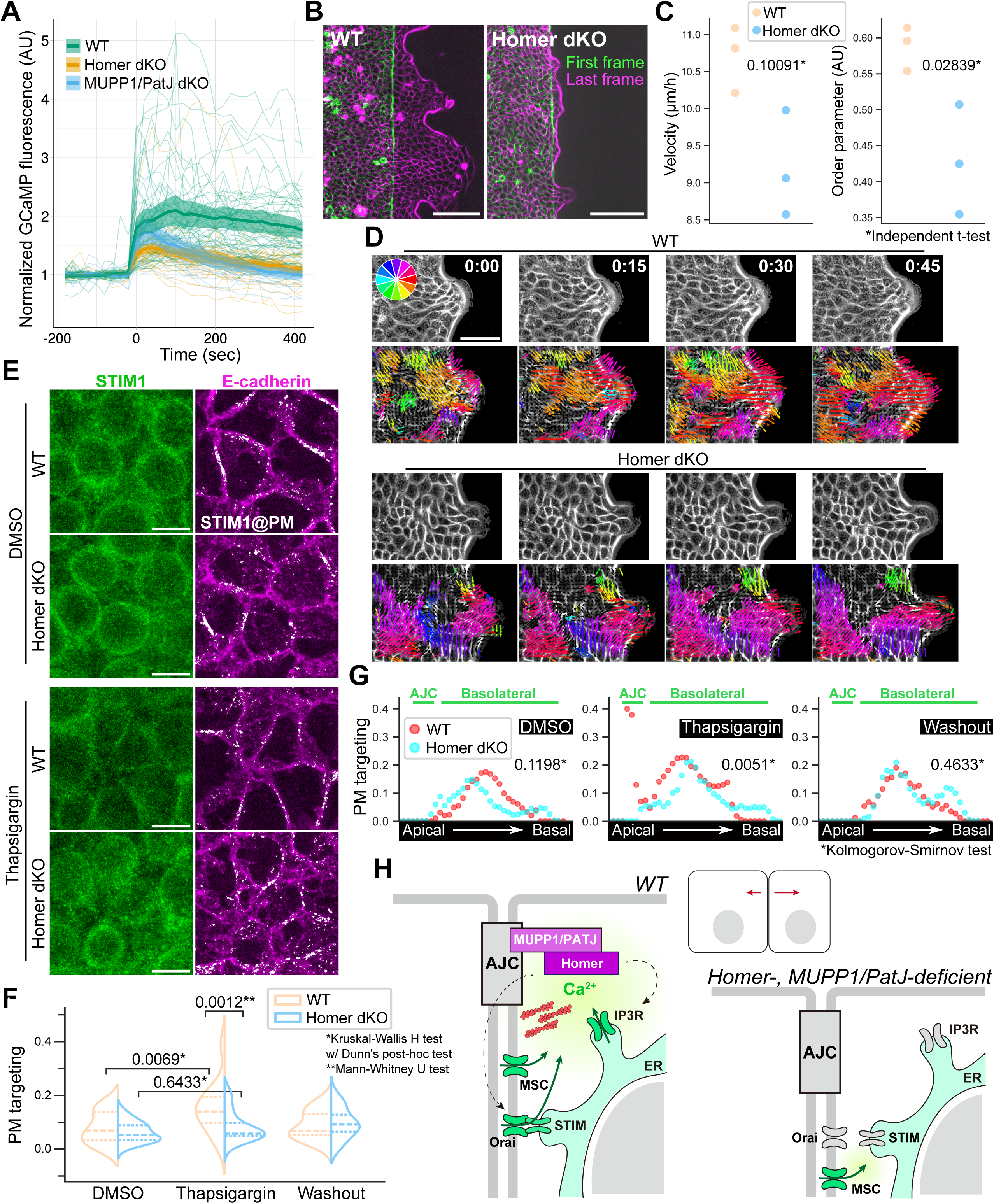
Signaling through Homer-MUPP1/PATJ controls mechanosensation in cultured epithelial cells. (A) GCaMP signals were normalized to pre-activation period. Individual signals (at least 50 cells per condition from three independent experiment) are plotted with the means and 95% confidence intervals in bold and shade, respectively. (B) Time overlay of migrating WT and Homer dKO MDCKII monolayers showing. Scale bar = 100 µm. Also see Movie S1. (C) Quantification of wound healing migration based on data obtained from PIV analysis. Velocity and order parameter, which describes multicellular coordination, are plotted. Each data point is the mean of one independent time lapse experiment. At least 1,000 directional vector pairs between consecutive frames were quantified over a total of 36 frames. (D) Phase contrast and PIV overlay of finger-like projections in the migrating monolayer. Scale bar = 50 µm. (E) Immunofluorescence images of WT and Homer dKO cells treated with thapsigargin stained for STIM1 and E-cadherin. Colocalized STIM1 is overlaid on the E-cadherin images. Also see Figure S5H. Scale bar = 10 µm. (F) Quantification of the degree of STIM1 colocalizing with E-cadherin. The rate of colocalization was calculated for individual z plane images from four images obtained over three independent experiments and collated. The total number of cells contained in the images were at least 50. Violin plots show the median with 25^th^ and 75^th^ percentiles. (G) Plot of the degree of STIM1 colocalization with E-cadherin of a representative confocal image stack. (H) Schematic of tension-induced amplification of calcium signaling controlled by Homer-MUPP1/PatJ. Abbreviations are as follows: AJC, apical junctional complex; MSC, mechanosensitive ion channel; ER, endoplasmic reticulum

To explore how this is translated to tissue-scale behavior, we employed the wound healing migration model of MDCKII epithelial cells, where cells transmit directional cues through directional propagation of intercellular tension^17^. As in EpH4 cells, MDCKII cells do not express Homer2. Therefore, we established Homer1,3 dKO cells and examined their migration by particle image velocimetry (PIV, Movie S1). Sheet migration of the Homer dKO monolayer was visibly impaired (Figures 3B). In contrast to the WT cell sheet that move toward the wound methodically, the Homer dKO monolayer remains stationary, even as the individual cells display roughly equivalent motility (Video S1 and Figure 3C). Particularly notable was the growth of collective behavior in the finger-like projections where WT cells steadily acquired directional coordination while in Homer dKO cells, swathes of cells moved independently of one another (Figures 3D). This impression was borne out by quantifying the order parameter^34^, which describes the direction of local motion relative to that of the migration edge. We observed that directional homogeneity expanded with time in the WT monolayer but was incongruous and volatile in Homer dKO cells, which is supported by a significant decrease in the order parameter in the Homer dKO cell sheet (Figures S4C and 3C). We conclude that Homer-MUPP1/PatJ mediates mechanosensation, which materializes as tissue-wide coordination during epithelial wound healing.

It is notable that tension application could induce an acute calcium response in cells lacking Homer and MUPP1/PatJ—though there was a marked decrease in its magnitude—which suggests that this this initial rise is not strictly dependent on our proposed mechanism. Mechanosensitive ion channels, such as Piezo and transient receptor potential (TRP) channels are implicated in various calcium responses; during *Drosophila* dorsal closure, pharmacological activation of mechanosensitive ion channels causes calcium influx and cell contraction while their inhibition or genetic ablation lead to defective actomyosin organization and failure of closure^20^. While Piezo channels are randomly localized throughout the plasma membrane and are diffusive, calcium flickers were shown to overlay Piezo channels in regions under high traction forces^35^. We found that TRPV4 was broadly distributed throughout the lateral membrane both in steady state EpH4 cells and in migrating MDCKII cells (Figures S5A and S5B); its localization was also unaltered in Homer knockout cells (Figures S5A).

Calcium inflow sensitizes the activation of IP3R and RyR to release the ER-stored calcium in a process known as calcium-induced calcium release. However, calcium has limited diffusion potential in the crowded milieu of the cytoplasm as it is quickly quenched, through either consumption and buffering by calcium binding proteins or uptake into intracellular stores^23^. Depletion of ER calcium then triggers amplifying, regenerative calcium signaling by engaging Store-operated calcium entry (SOCE), in which the oligomerization of the ER calcium sensor STIM1 activates the plasma membrane (PM) calcium channel Orai1 to induce calcium influx from the extracellular space. Additionally, STIM1 can amplify local calcium signaling by recruiting the TRPC family of TRP cation channel superfamily^36^. Homer1 is reportedly found in complex with Orai1 and STIM1 and positively regulates SOCE^6,14^. Suppression of SOCE and mechanosensitive ion channels by inhibitor treatment, and SOCE specifically by STIM1 knockout, diminished AJC actin enrichment (Figures S5C-S5E). We then activated SOCE by treating cells with thapsigargin to empty the ER store in order to examine the role of Homer in STIM1 localization. In untreated cells, STIM1 was prominent perinuclearly and in the vicinity of the PM along the lateral membrane (Figure 3E and S5F). Thapsigargin treatment significantly enhanced its targeting to the PM in WT but not Homer dKO cells (Figure 3F). This was due to the appearance of AJC-targeted STIM1 specifically in WT cells since lateral STIM1 was not altered by Homer depletion (Figure 3G). Altogether, these data implicate SOCE as the target of Homer-MUPP1/PatJ. We therefore posit that calcium entry via mechanosensitive ion channels trigger sustained calcium input at AJC and that Homer-MUPP1/PatJ are necessary to amplify this process by linking the initial calcium influx via mechanosensitive ion channels to SOCE and the ER calcium store (Figure 3H).

### Homer3 is required for neural tube closure during Xenopus embryogenesis

Notably, mutation in MUPP1 in humans is linked to nonsyndromic congenital hydrocephalus, which occurs as a result of defective neural tube closure^37,38^, a calcium-dependent process in *Xenopus*^22,39^. This is possibly due to the requirement of MUPP1 for apical constriction of neuroepithelial cells and subsequent neural plate bending that precedes the formation of the neural folds, which converge to form the neural tube^40^. In addition, the apical constriction of neuroepithelial cells during neural tube closure requires asynchronous transient elevations in intracellular calcium ion levels^22,39^, and knockout mice lacking Secretory Pathway Calcium ATPase 1 (SPCA1), which is involved in calcium homeostasis, exhibit phenotypes indicative of defective neural tube closure^41^. We thus turned to the *Xenopus* model to examine the physiological function of signaling through Homer. Analysis of the expression pattern of Homer isoforms in relation to developmental stages at the Xenbase database revealed that Homer3 (*homer3.L*) is specifically upregulated during neurulation (stage 12-20, Figure 4A)^42,43^. Two independent morpholinos designed to block the splicing of Homer3 pre-mRNA (MO1 and MO2) were unilaterally injected with the EGFP marker mRNA at the 4-cell stage and embryos were sampled at stage 18/19 to assess neurulation defects (Figure 4B). Disruption of Homer3 expression delayed neural tube closure in 91% of MO1– and 100% of MO2-injected embryos, whereas no abnormality was observed in those injected with the control morpholino (Standard MO, Figures 4C and 4D). The apical domain of the superficial cells in the Homer3 MO-injected side were enlarged compared to the uninjected side and displayed less junctional actin enrichment, indicating a defect in apical constriction (Figures 4E and 4F). This observation agrees with the role of MUPP1 in neurulation where failure of apical constriction was reported to cause the delay in neural tube closure^40^. Thus, we surmise that Homer cooperates with MUPP1 to drive tissue remodeling during neurulation.

**Figure 4.**
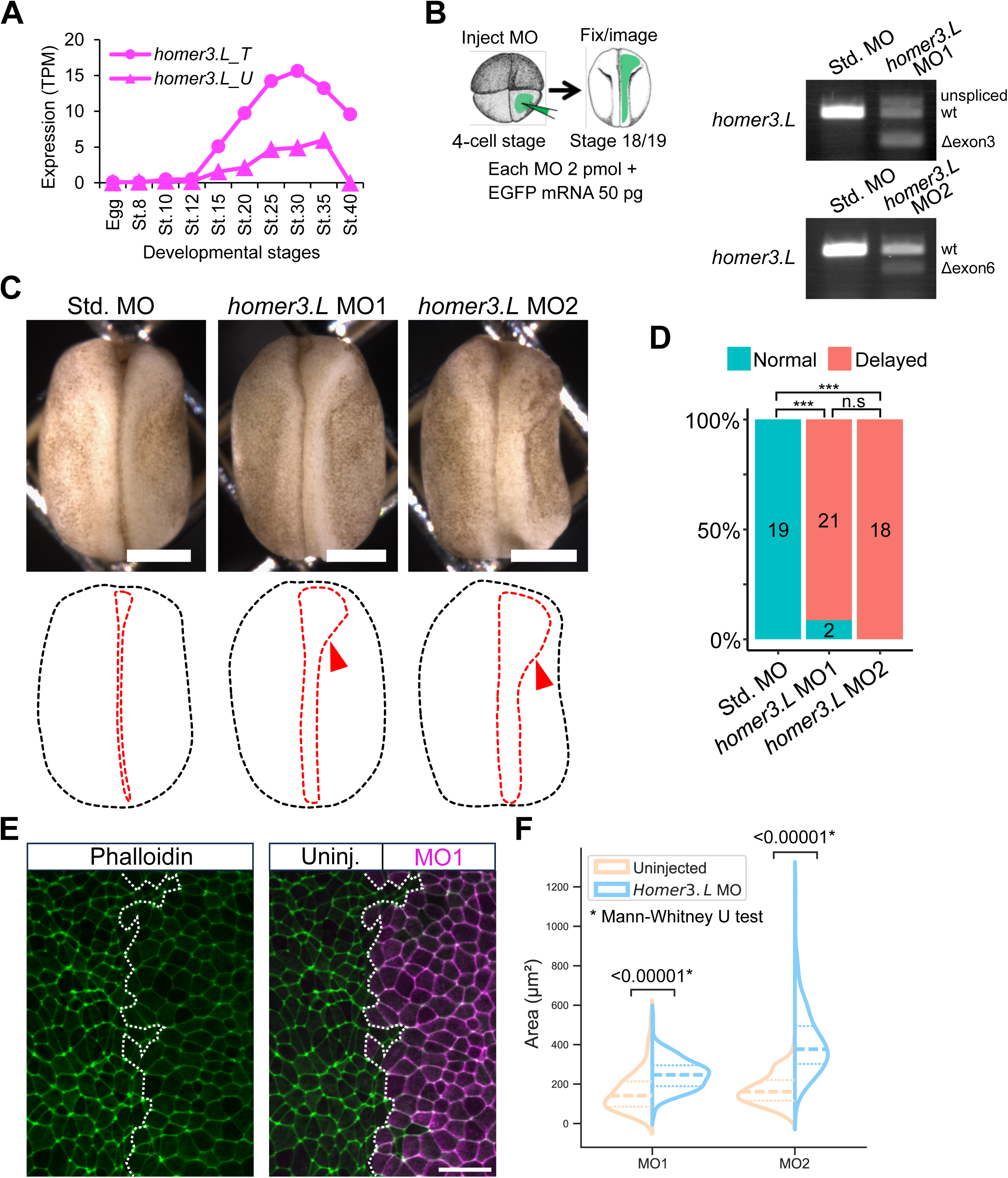
Homer3 is required for neural tube closure during *Xenopus* embryogenesis. (A) Expression pattern of *homer3.L* in whole embryos at various developmental stages. Graph label suffixes “_T” and “_U” indicate the independent RNA-seq data sets “Taira201203” and “Ueno201210”, respectively. (B) Top: Experiment schematic. Either Control (Std. MO) or targeted morpholino (*homer3.L* MO1 and MO2), together with the EGFP marker mRNA, was injected into the right-dorsal blastomere of the 4-cell-stage embryo. Bottom: RT-PCR of homer3.L MO-injected embryos. Longer and shorter PCR products compared with wt indicate unspliced and exon-skipped *homer3.L* mRNAs, respectively. The decreased homer3.L expression suggests nonsense-mediated decay of the transcript. (C) and (D) Representative micrographs of the embryos (C) showing delayed neural tube closure in *homer3.L* MO1– and MO2-injected embryos (red arrowheads). The dotted black and red lines indicate the outline of the embryo and neural tissue, respectively. Scale bar = 500 µm. The results are quantified in (D). *P*-values were calculated using pairwise Fisher’s exact test with Bonferroni correction (\*\*\**P* < 0.001; n.s., not significant). (F) Phalloidin staining of a representative stageL15–16 embryo showing uninjected and morpholino-injected hemispheres. Scale bar = 50 µm. (G) Quantification of (E) showing cell area. Three fields of view per embryo were imaged and quantified. Three embryos and five embryos were analyzed for MO1 and MO2 respectively. At least 200 cells were quantified per image and collated.

These results suggest that the function of Homer-MUPP1/PatJ in transducing mechanical information is conserved in cultured epithelial cells of disparate lineage, mammary (EpH4) and kidney (MDCKII), and in developing tissue (*Xenopus*). However, we acknowledge that further study is required to establish conservation through *Xenopus* of the specific molecular mechanism, i.e. the functional interaction between Homer and MUPP1 and calcium signaling.

In summary, our study provides new insights into tension sensing at contractile interfaces in epithelial tissue, laying out the groundwork for understanding how local calcium dynamics can regulate tissue-scale changes. We show that there is an adaptive, tension-sensitive pool of actin at AJC, which is potentially regulated by a distinct calcium signature composed of signal variance and amplitude. Calcium input at AJC requires amplification of the signal mediated by Homer and MUPP1/PatJ scaffolding proteins to overcome calcium quenching by intracellular buffers. The Homer-calcium signaling module enables intercellular tension sensing, which informs collective tissue behavior such as wound healing migration in a tissue culture model and neurulation in *Xenopus* embryogenesis. Regarding the latter, the observed effect on cell morphology, i.e. the enlarged apical surface area of Homer-deficient cells at the dorsal midline, leaves open the possibility that depressed baseline contractility due to Homer depletion is at its root, since apical constriction is widely considered the primary driver of tissue folding^44^.

However, neural tube closure involves a complex interplay between multiple cell behaviors besides apical constriction, such as cell intercalation and directed protrusion^45–47^. One recent model argues that heterogeneity of cell populations enables constricting cells to instruct the alignment of non-constricting cells and cause folding^48^. Another model proposed that PCP-dependent anisotropic junction shrinking is sufficient to generate cellular forces that would cause neural plate bending^45^. Tension sensing through Homer-calcium signaling could play a role in either case to coordinate tissue-scale response and merit investigation, perhaps through targeted downregulation of the module in specific cell populations. The prevailing view on the etiology of neural tube defects, including hydrocephalus, suggests that it arises from a dysfunction in the ciliated ependymal cells lining the ventricles, either due to impaired absorption of excess cerebrospinal fluid or overproduction of cerebrospinal fluid^49,50^. Future research should focus on understanding how the tension-dependent calcium influx mechanism, involving MUPP1 and Homer as highlighted in this study, contributes to the pathogenesis of these conditions.

### Materials and methods

#### Regents, plasmids and morpholinos

EpH4, MDCKII and HEK293 cells were cultured in 10% (vol/vol) FCS/DMEM. Knockout epithelial cells were generated by CRISPR/Cas9 gene editing according to Zhang laboratory protocols (F. Zhang, Massachusetts Institute of Technology, Cambridge, MA; https://www.addgene.org/crispr/zhang/) using the following target sequences: MUPP1 (mouse), 5′-GCTTCCCGCTGTGTCCCACA-3′; PATJ (mouse),

5′-GTAAGGATCTGGTTGAAGAG-3′; Homer1 (mouse),

5′-GCAAACACTGTTTATGGACT-3′; Homer3 (mouse),

5′-GTAAGTGCGTGCTTGCCGGC-3′; Homer1 (dog), 5’-

GTAACTGCATGCTTGCTGGT-3’: Homer3 (dog), 5’-

GATTCGGTAGACATTTCGGG-3’; STIM1 (mouse), 5’-

GAATACAGGAGCTAGCTCCG-3’.

We used the following antibodies for immunofluorescence microscopy and immunoblotting: mouse anti-MPDZ mAb (611558, BD Transduction Laboratories), rabbit anti-INADL/PATJ pAb (LS-C410011-100, LSBio), rat ECCD-2 mAb (Takara Bio), mouse anti-claudin-4 mAb (32-9400, Life technologies), rabbit anti-Homer1 pAb (GTX103278, GeneTex), rabbit anti-Homer3 (NBP2-32607) pAb (Novus Biologicals), rabbit anti-STIM1 mAb (D88E10, Cell Signaling Technology), rabbit anti-TRPV4 pAb (ab39260, abcam) and mouse anti-DYKDDDDK mAb (1E6, FUJIFILM Wako Chemicals). Rat anti-GFP mAb (JFP-J5), mouse anti-α-tubulin mAb (12G10), mouse anti-ZO-1 mAb (T8754) and mouse anti-Myc mAb (9E10) were produced from hybridoma in our lab. The secondary antibodies for immunofluorescence were purchased from Jackson ImmunoResearch and they were as follows: anti-rat IgG antibodies (Cy2-conjugated, 712-225-150 and Cy3-conjugated, 712-165-150); anti-mouse IgG antibodies (Cy2-conjugated, 715-545-151 and Cy3-conjugated, 711-165-150); and anti-rabbit-IgG antibodies (Cy2-conjugated, 711-225-152 and Cy3-conjugated, 711-165-152). The HRP-conjugated secondary antibodies for immunoblotting were: anti-rat IgG antibody (HAF005, R&D Systems); anti-mouse-IgG antibody (A90-516P, Bethyl Laboratories); and anti-rabbit IgG antibody (4030-05, Southern Biotech). Alexa Fluor 488–phalloidin was used to detect F-actin(A12379; Invitrogen).

The sequence of *X. laevis homer3.L* and RNA-seq data (GSE73430, NCBI GEO) were analyzed using Xenbase^43^ and *Xenopus* genome database (http://viewer.shigen.info/xenopus/). Splicing-blocking antisense morpholino oligonucleotides (MOs) that target the splice donor sites of exon 3 (MO1, 5’-AACAAGATATCATCCCTCACCTTG-3’), and exon 6 (MO2, 5’-AAAAGAGCACAACTTACCCTTCGGA-3’) of *homer3.L* were designed and synthesized by GeneTools. A pre-designed standard MO (5’-CCTCTTACCTCAGTTACAATTTATA-3’) was used as a negative control. The splicing defects of *homer3.L* mRNA were examined by RT-PCR analysis using RNeasy Mini Kit (Qiagen) and PrimeScript II High Fidelity RT-PCR Kit (Takara Bio). The sequences of PCR primers used are as follows: MO1 forward,

5’-CGCAGGCTTTCATCCACAAATG-3’; MO1 reverse,

5’-TGAAGCGAAGCCCAATCCATAC-3’; MO2 forward,

5’-GCAAGGGAGAGATCTCAGGACAAG-3’; MO2 reverse,

5’-TAGCATTTCCGCCTCCTCCTGATA-3’.

#### Live cell imaging

Cells were cultured in 10% (vol/vol) FCS/Leibovitz’s L-15 Medium (1415064; Thermo Fisher Scientific) to support cell growth in atmospheric CO_2_ concentration. Images were acquired using a confocal microscope (LSM900; Carl Zeiss MicroImaging) equipped with a Plan-APO (63×/1.40 NA, oil immersion) objective and controlled by Zen 2012 software (Carl Zeiss Microimaging); cells were imaged on a heating stage set to 37 °C. Cells for wound healing migration were seeded in Culture-Insert 2 Well (ibidi) and cultured to confluence overnight prior to insert removal. Live cell imaging was performed using BZ-X710 all-in-one fluorescence microscope equipped with a time-lapse module, Plan Fluor 20x/o.45 NA objective (Nikon) and a heating stage set to 37 °C. Phase contrast images were captured with the on-board CCD camera. OptoRhoGEF was activated by irradiation of the entire sample with blue light from an LED light source at 100% power. Cells were treated with 1 µM BAPTA-AM (DOJINDO LABORATORIES) or with 1 µM A23187 (Sigma-Aldrich) after a period of imaging at steady state.

#### Fluorescence microscopy

Cells were exposed to the following inhibitors for 30 min prior to fixation: BAPTA-AM (DOJINDO LABORATORIES) was used at 1 µM, 2-Aminoethyl diphenylborinate (2-APB, Tocris Bioscience) at 50 µM, Xestospongin C (FUJIFILM Wako Chemicals) at 10 µM, SKF96365 (Tocris Bioscience) at 10 µM A23187 (C7522), AnCoA4 (Calbiochem) at 50 µM and thapsigargin (FUJIFILM Wako Chemicals) at 1 µM. Cells were fixed either with 100% methanol at –20℃ for 20 min, or with 3% formalin/PBS for 15 min at RT. Formalin-fixed cells were permeabilized with 0.4% Triton X-100/PBS for 5 min. Cells were blocked with 1% BSA/PBS for 1 h at RT and reacted with primary antibodies for 1 h and secondary antibodies for 30 min at RT. Antibodies were diluted in blocking buffer. Anti-PATJ pAb was prepared in Can Get Signal Immunostain Solution B (NKB-601; TOYOBO). Image were acquired using the same confocal microscopy system as in live cell imaging.

#### Xenopus embryo culture, microinjections, and morphological analyses

Experiments with *X. laevis* embryos were performed as previously described^22^. Briefly, embryos were raised by in vitro fertilization^51^. 2 pmol MO was injected into the right dorsal blastomere (prospective neuroectoderm) of 4-cell-stage embryos together with 50 pg EGFP mRNA or 100 pg membrane-targeted mRFP mRNA. The injected embryos were cultured in 3% Ficoll/0.3× MMR to stage 9 and then cultured in 0.3× MMR^51^ until appropriate stages^52^. For observation, embryos were fixed for 2 h with modified MEMFA (0.1 M MOPS [pH 7.4], 2 mM EGTA, 1 mM MgSO4, 3.7% formaldehyde, and 1% glutaraldehyde). Bright-field images were acquired using an Axio Zoom.V16 zoom microscope (Carl Zeiss Microimaging) and analyzed using Zen (Carl Zeiss Microimaging). For F-actin visualization, embryos were fixed for 2 h with MEMFA and stained with Alexa Fluor 488–phalloidin (A12379; Invitrogen). Confocal images were acquired using a spinning-disc confocal unit (CSU-W1; Yokogawa) with an EMCCD camera (iXon Ultra 888; Andor) on an inverted microscope (ECLIPSE Ti2; Nikon) with a 20× (Plan Flour 20×/0.50; Nikon) or 60× (Plan Apo VC 60×A WI/1.20; Nikon) objective and analyzed using ImageJ/Fiji software. Data plots and statistical analyses were performed using the R software (R Core Team), Python and Excel (Microsoft).

#### Image processing, quantification and statistical analysis

Images were prepared for quantification using ImageJ/Fiji and a Napari package (github.com/haesleinhuepf/devbio-napari).

Kymographs were produced in ImageJ from single AJC plane time lapse images using 10 pixel-width lines drawn perpendicular to the GCaMP-CAAX signal and masked to eliminate background. GCaMP-CAAX signals were mean intensities of the line width and displacement data were derived from maximum signal intensities at each time point.

The following python packages were used: numpy, pandas, statsmodels and scipy for organizing, sorting and processing (normalization, smoothing, peak/trough finding) to automatically determine analysis windows based on displacement and extract data for various parameters; statsmodels for OLS analysis; matplotlib and seaborn for presentation. Any analysis window with a single peak/trough was automatically thrown out.

Cross-correlation was analyzed between junction deformation and calcium signal using the python packages numpy, pandas, matplotlib, statsmodels and scipy. Time series were normalized, smoothed using the LOWESS model and tested for stationarity using the Augmented Dickey-Fuller test; non-stationary time series were detrended before calculating cross-correlation. Statististically significant lags exceeding the 95% confidence interval threshold for 7 of 12 junctions analyzed. For control, cross-correlation between random pairs of junction deformation and calcium signal time series were calculated until 7 significant lags matching the result of the experimental dataset was obtained. We then compared the means and data variance between the experimental and randomized control datasets using Mann-Whitney and Levene’s tests, respectively.

Overall cell deformation was assessed by quantifying the frame-to-frame change in cell area normalized for the mean area of the cell across the time lapse. Areas were obtained by segmentation using Cellpose3^53,54^. The python packages numpy, pandas, skimage, scipy, matplotlib and seaborn were used to process, measure and analyze the masks and plot the data. Processed masks were visually inspected to ensure that the same cell was tracked across all time frames after analysis. Change point detection was performed by the PELT method^55^ in the ruptures package. Quantification of actin enrichment at AJC was performed essentially as described^15^. Confocal stacks of ZO-1 (tight junction, a proxy for AJC) and E-cadherin (adherens junction) images were segmented using the Convpaint^56^ plugin in Napari to create AJC and lateral masks for phalloidin image processing. STIM1 and E-cadherin images were likewise segmented for analysis of colocalization, defined as the intersection of segmented label images.

The PlotTwist web app (https://huygens.science.uva.nl/PlotTwist)^57^ was used for cluster analysis based on Manhattan distance and complete linkage method. Clustering was verified with the Calinski Harabasz index. PIV analysis was performed using the ImageJ plugin TPIV (https://signaling.riken.jp/tools/imagej-plugins/490/) after masking for the cell-free area. Order parameter was calculated as described^34^.

Normality of data distribution was assessed by the Shapiro-Wilk test and means were compared by either Independent t-test or Mann-Whitney U test as appropriate. Either ANOVA with Tukey’s or Kruskal-Wallis H test with Dunn’s post-hoc test was chosen as appropriate for multiple comparison. Kolmogorov-Smirnov test and Levene’s test were used to assess the variability in distribution and variability in the dataset. All quantifications are representative of at least three independent experiments. Statistical tests and significance are noted in the figures.

#### Immunoblotting

Cells were lysed in SDS sample buffer. SDS samples were resolved by SDS-PAGE, transferred to nitrocellulose membranes, blocked with 5% (wt/vol) skim milk/0.1% (vol/vol) Tween 20/TBS and sequentially reacted with primary antibodies diluted for 1 h and with HRP-conjugated secondary antibodies for 30 min, both prepared in blocking buffer. Chemiluminescence was imaged using the LAS4000mini imaging system (Fujifilm).

#### FLAG immunoprecipitation

Cells were washed with ice-cold PBS then lysed with cell lysis buffer (20 mM Tris-HCl [pH 7.5], 150 mM NaCl, 1% Triton X-100) containing the following protease inhibitors: (leupeptin (10 μg/ml, 334-40414, Wako Pure Chemical Industries); aprotinin (2 μg/ml,10236624001 Roche); and amidinobenzylsulfonyl fluoride (50 μM, 015-26333, Wako Pure Chemical Industries). Lysates were centrifuged to remove insoluble content and clarified lysates were incubated with anti-DYKDDDDK tag antibody beads (018-22783; FUJIFILM Wako Pure Chemical) overnight. Bound proteins were eluted with SDS sample buffer after extensive washing of the beads.

## Acknowledgments

We thank all members of the Ikenouchi laboratory (Department of Biochemistry, Graduate School of Medical Sciences, Kyushu University) for helpful discussions. We also thank Dr. Alexander Ludwig (Nanyang Technological University) for engaging in insightful discussions. Additionally, we acknowledge the National Bio Resource Project (NBRP) of MEXT for providing the *Xenopus* genome database.

## Author contributions

K.M. performed most of the experiments, analyzed the data and wrote the paper. M.S. designed and performed in vivo analyses using *Xenopus laevis* embryos. Y.C. and R.F. performed some experiments. J.I. designed research and wrote the paper.

