## Appendix 01 for "A steady state pool of calcium-dependent actin is maintained by Homer and controls epithelial mechanosensation"

**This PDF file includes:**

Figures S1 to S5
Legend for Movie S1

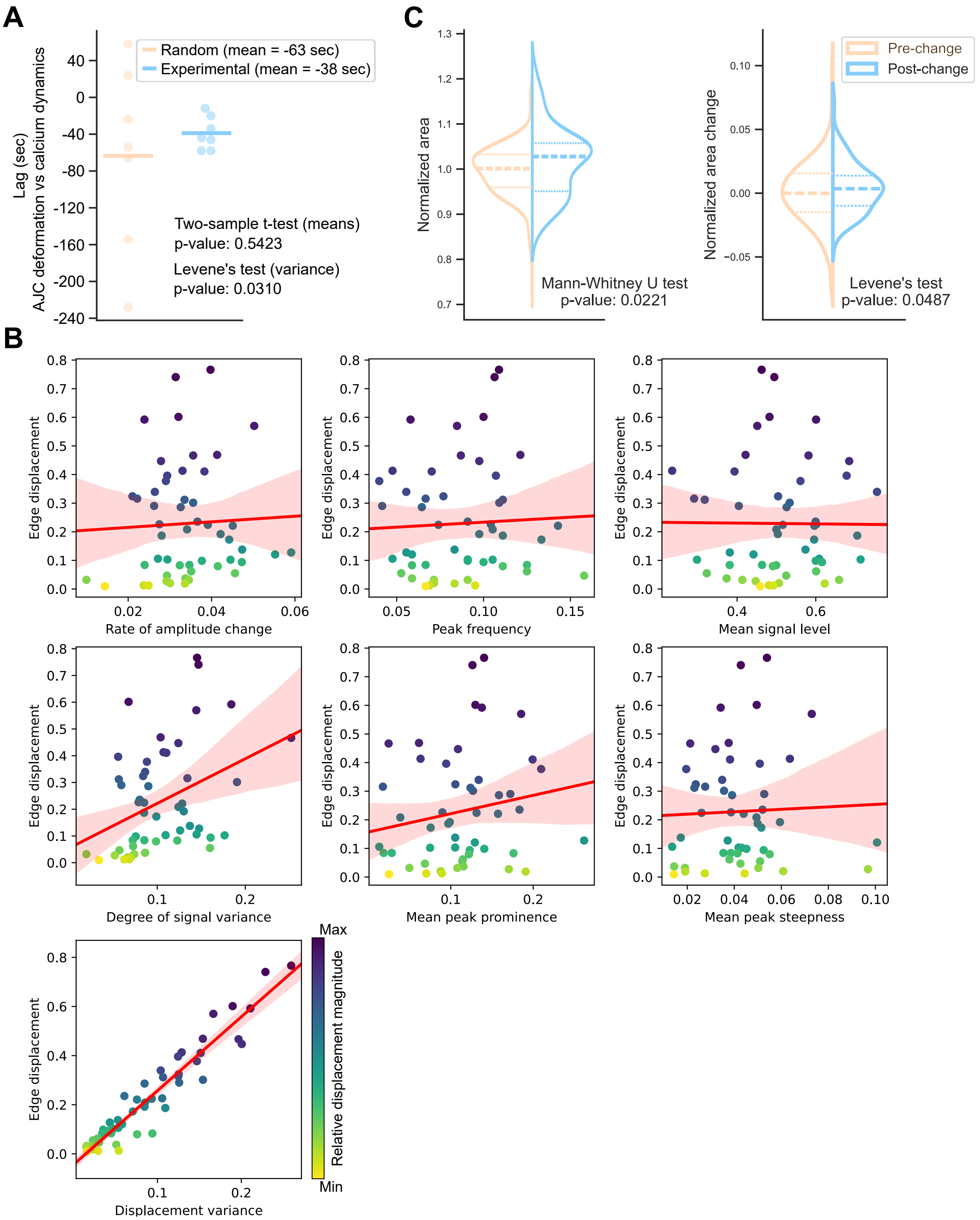


**Fig. S1. Relationship between calcium dynamics and cell morphological changes**
(A) Plot showing the lags derived from cross correlation analysis between junction movement and calcium dynamics time series. Statistically significant lags exceeding the 95% confidence interval threshold were obtained for 7 of 12 junctions analyzed. Control data set was derived by performing cross correlation analysis between random pairs of junction movement and calcium dynamics time series until we obtained 7 significant lags. Results of the statistical analyses for means and variance comparison are indicated.

(B) Scatter plots of calcium dynamics variables plotted against the corresponding magnitude of edge displacement overlaid with regression lines.
(C) Normalized areas and rate of area change in Figure 1E were compared before and after the change-point. Violin plot shows the median with 25th and 75th percentiles

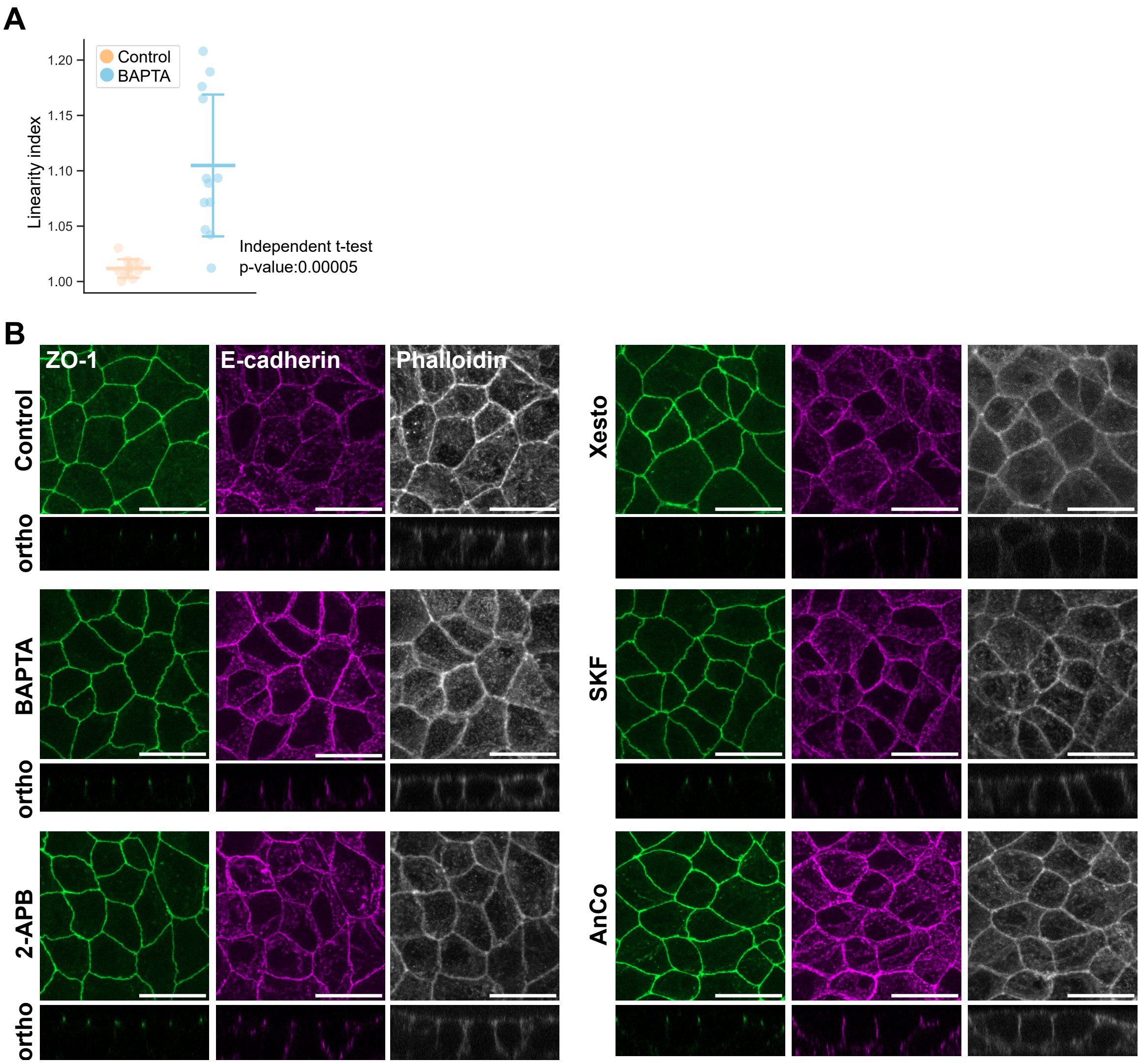


**Fig. S2. Effect of cytoplasmic calcium inhibition on AJC and actin structures (related to Figures 1 and S5)**
(A) Quantification of junction linearity of images represented in Figure 1F. Linearity index is defined as the ratio of the actual junction length over the hypothetical straight line. Twelve junctions from four individual cells imaged over three independent experiments are quantified. (B) Immunofluorescence images of WT cells treated with the indicated inhibitors stained for ZO-1, E-cadherin and phalloidin. BAPTA-AM is a cell permeable calcium chelator, 2-APB inhibits IP3R and numerous TRP channels, Xestospongin C targets IP3R. SKF96365 targets TRP channels and SOCE and AnCoA4 primarily targets SOCE.

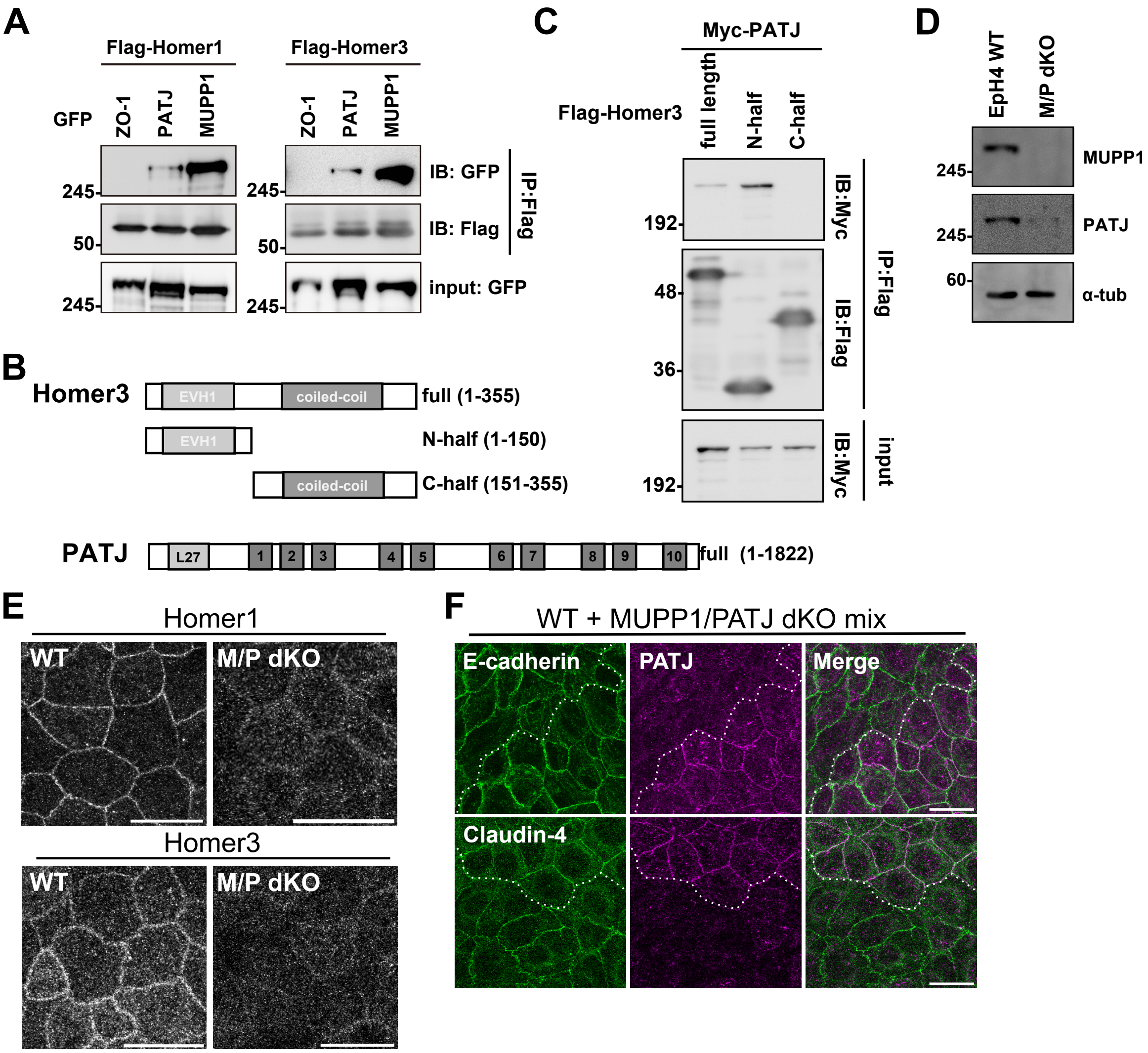


**Fig. S3. Homer-PATJ interaction in detail (related to Figure 2)**

(A) Immunoblots of FLAG immunoprecipitates probed with the indicated antibodies. FLAG-tagged Homer 1 or Homer 3 were co-expressed with one of GFP-tagged ZO-1 (negative control), MUPP1, or PATJ.

(B) Domain structures and the analyzed fragments of Homer3 and PATJ.

(C) Immunoblots of FLAG immunoprecipitates probed with the indicated antibodies. FLAG-tagged Homer 3 fragments were co-expressed with myc-tagged PatJ.

(D) Whole-cell lysates of WT andMUPP1/PATJ dKO cells were immunoblotted with the indicated antibodies.

(E) Immunofluorescence images of WT and MUPP1/PATJ dKO cells stained for Homer1 or Homer3. Scale bar = 20 μm.

(F) Immunofluorescence images of a co-culture of WT and MUPP1/PATJ dKO cells stained for either Claudin-4 or E-cadherin. Scale bar = 20 μm.

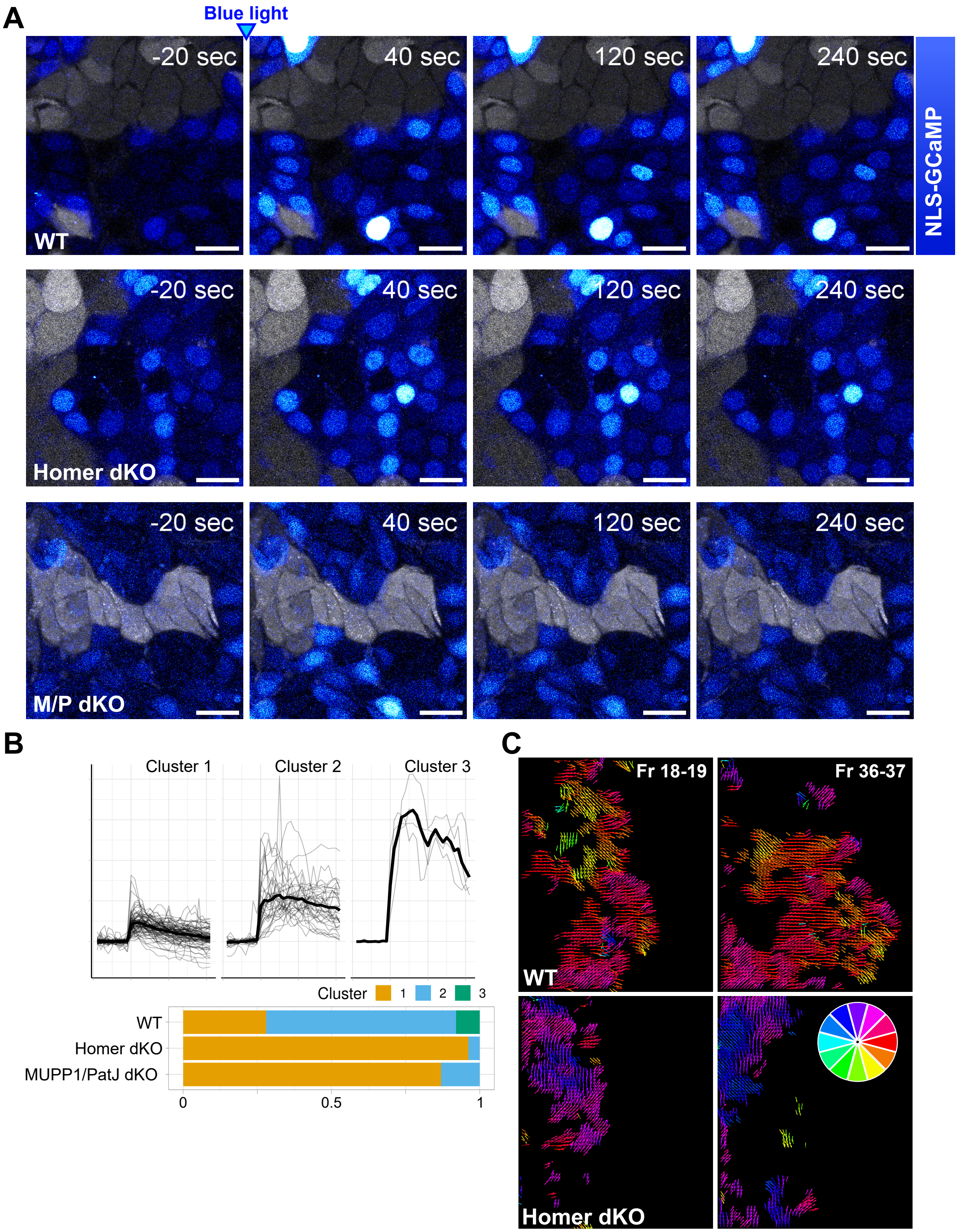


**Fig. S4. Role of Homer-MUPP1/PatJ in regulating tissue-scale behavior requiring mechanosensation (related to Figure 3)**
(A) Time lapse images of a co-culture of OptoRhoGEF-expressing cells and GCaMP-NLS-expressing WT, Homer dKO and MUPP1/PatJ dKO cells. Scale bar = 20 µm.
(B) GCaMP signals from Figure 3A were clustered by Manhattan distance with complete linkage. AT least 50 WT cells, 52 Homer dKO cells and 43 MUPP1/PatJ dKO cells were quantified over three independent experiments

(C) PIV images from Figure 3B and Movie S1. Directional vectors were colored as shown in the color wheel.

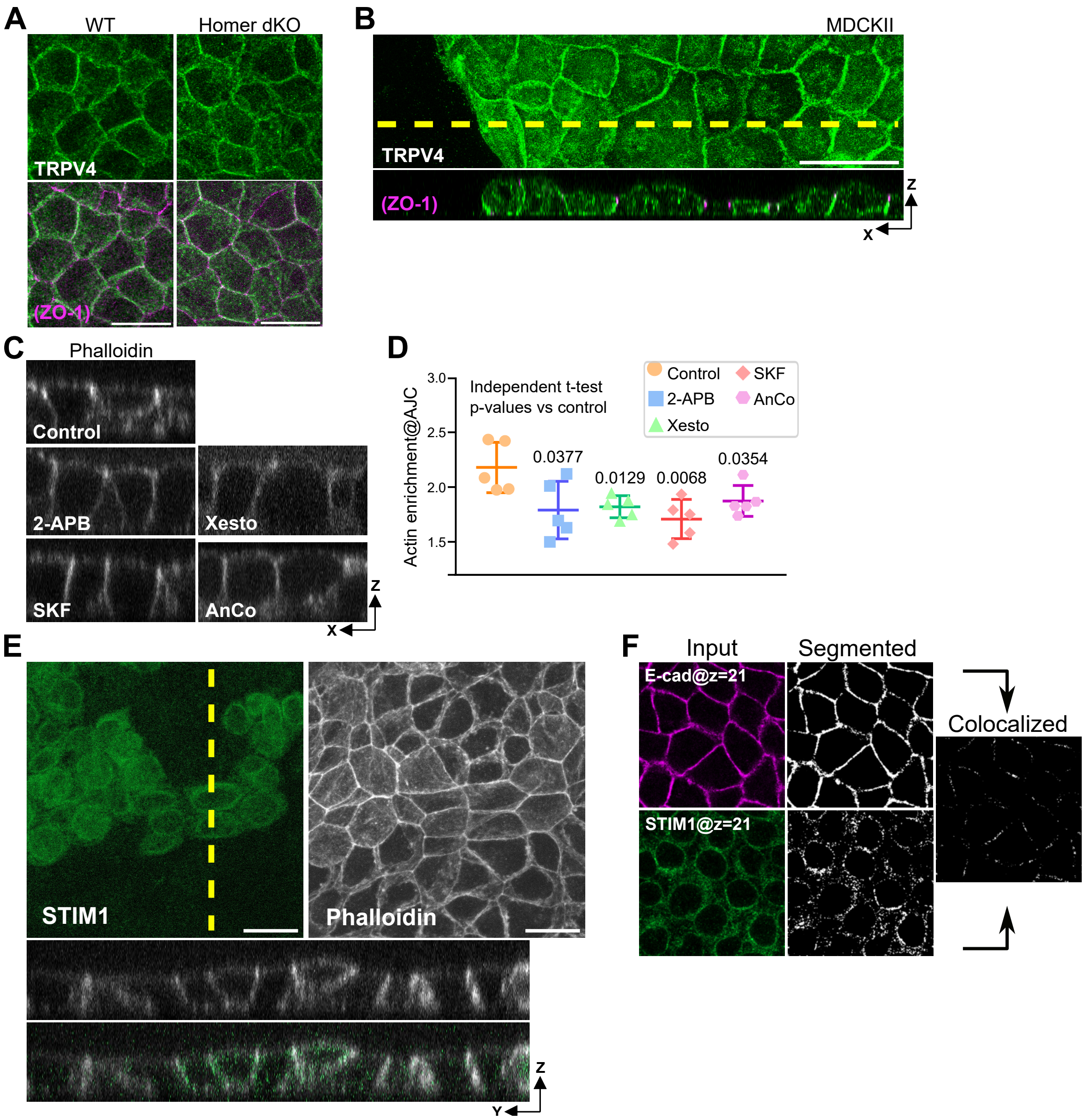


**Fig. S5. Involvement of SOCE in mediating Homer-dependent regenerative calcium input**
(A) Immunofluorescence images of WT and Homer dKO EpH4 cells stained for TRPV4 and ZO-1. Colocalized STIM1 is overlaid on the E-cadherin images. Also see Figure S5H.
(B) Immunofluorescence image of a migrating MDCKII monolayer at the wound edge stained for TRPV4 and ZO-1.
(C) Phalloidin-stained images of WT cells treated with the indicated inhibitors.
(D) Quantification of (C). ZO-1 and E-cadherin staining were used to define AJC and lateral regions to quantify actin enrichment at AJC. Mean and standard deviation are indicated. Individual dots are means of one field of view and five independent images obtained over three independent experiments were analyzed.
(E) Immunofluorescence images of WT and STIM1 KO EpH4 cells stained for STIM1 and phalloidin.
(F) Workflow of the colocalization analysis.
Scale bars = 20 μm.

**Movie S1. Time lapse and corresponding PIV of migrating MDCKII WT and Homer dKO monolayers**
Phase contrast and PIV movies are presented together. Scale bar = 200 μM.
